# A genome-wide screen in mice to identify cell-extrinsic regulators of pulmonary metastatic colonisation

**DOI:** 10.1101/2020.02.10.941401

**Authors:** Louise van der Weyden, Agnieszka Swiatkowska, Vivek Iyer, Anneliese O. Speak, David J. Adams

## Abstract

Metastatic colonisation, whereby a disseminated tumour cell is able to survive and proliferate at a secondary site, involves both tumour cell-intrinsic and -extrinsic factors. To identify tumour cell-extrinsic (microenvironmental) factors that regulate the ability of metastatic tumour cells to effectively colonise a tissue, we performed a genome-wide screen utilising the experimental metastasis assay on mutant mice. Mutant and wildtype (control) mice were tail vein-dosed with murine metastatic melanoma B16-F10 cells and 10 days later the number of pulmonary metastatic colonies were counted. Of the 1,300 genes/genetic locations (1,344 alleles) assessed in the screen 34 genes were determined to significantly regulate pulmonary metastatic colonisation (15 increased and 19 decreased; *P*<0.005 and genotype effect <-60 or >+60). Whilst several of these genes have known roles in immune system regulation (*Bach2, Cyba, Cybb, Cybc1, Id2, Igh-6, Irf1, Irf7, Ncf1, Ncf2, Ncf4* and *Pik3cg*) most are involved in a disparate range of biological processes, ranging from ubiquitination (*Herc1*) to diphthamide synthesis (*Dph6*) to Rho GTPase-activation (*Arhgap30* and *Fgd4*), with no previous reports of a role in the regulation of metastasis. Thus, we have identified numerous novel regulators of pulmonary metastatic colonisation, which may represent potential therapeutic targets.

## INTRODUCTION

Metastasis is the spread of cancer cells to a secondary site within the body, and is the leading cause of death for cancer patients. This multi-step process requires tumour cells to survive in the circulation (‘circulating tumour cells’, CTCs), leave the circulation (‘extravasate’) at a distant site, survive at this new site (‘disseminated tumour cells’, DTCs) and eventually proliferate, thus becoming a clinical problem. Circulating tumour cells can be found in the patient’s blood at the time of diagnosis, and thus it is highly likely that metastasis has already occurred, especially since surgical excision of a primary tumour does not always prevent metastasis (Talmadge and Fidler 2010). Indeed, studies have demonstrated that the early steps of the metastatic process are relatively efficient, with the post-extravasation regulation of tumour growth (‘colonisation’) being critical in determining metastatic outcome (Chambers et al. 2001). Thus, the prevention of metastasis is unlikely to be of therapeutic benefit, and a focus on preventing the survival of CTCs and/or growth of DTCs would seem a more feasible approach (Fidler and Kripke 2015).

The survival and growth of metastatic cells involves contributions from both tumour cell intrinsic factors and tumour cell extrinsic factors such as the microenvironment (‘host’), which includes stromal cells and the immune system (Quail and Joyce 2013). In recent years there has been a revolution in our understanding of the role that host factors, such as the immune system, stroma and vasculature play in the process of cancer progression. This is evidenced by the development of agents, such as checkpoint inhibitors, that provoke the immune system to identify and eliminate cancer cells. Importantly, studies in mice have made a significant contribution to these breakthroughs, such as with the clinically relevant PD-1 (Zago et al. 2016)and CTLA4 receptors (Leach et al. 1996), which were first identified and functionally characterised using mouse model systems. For this reason we sort to develop a genetic screen to identify new genes as tumour cell extrinsic regulators of metastatic colonisation.

In designing our screen we aimed to, where possible, unbiasedly screen mouse mutants to identify new genes involved in colonization of the lung, a common site of metastatic seeding for many tumour types. To this end, we used the ‘experimental metastasis assay’, which we have previously demonstrated is a sensitive, robust, and high-throughput method for *in vivo* quantification of the ability of metastatic tumour cells to colonize a secondary organ (Speak et al. 2017), to screen mutant mouse lines generated as part of the International Mouse Phenotyping Consortium (Meehan et al. 2017). In this paper we describe a collection of mutants identified over 7 years of screening (1,300 mutant mouse lines). This study reveals previously unappreciated pathways and processes that regulate this biology.

## MATERIALS & METHODS

### Mice

The mutant mice were generated as part of the International Mouse Phenotyping Consortium (Meehan et al. 2017), using either targeted embryonic stem cell clones obtained from the European Conditional Mouse Mutagenesis (EUCOMM) Programme/Knockout Mouse Project (KOMP)-CSD collection or EUCOMMTools or CRISPR/Cas9 technology to either genetrap the target transcript or disrupt a critical exon or to create a point mutation, as detailed previously (van der Weyden et al. 2017b). The vast majority of lines (>98%) were on the C57BL/6 background, with other strain backgrounds including 129 and FVB (strain-matched control mice were always used for each mutant line). The care and use of all mice in this study were in accordance with the Home Office guidelines of the UK and procedures were performed under a UK Home Office Project licence (PPL 80/2562 or P6B8058B0), which was reviewed and approved by the Sanger Institute’s Animal Welfare and Ethical Review Body. All mice were housed in individually ventilated cages in a specific pathogen free environment. The diet, cage conditions and room conditions of the mice were as previously reported (van der Weyden et al. 2017a).

### Cells for tail vein injection

The B16-F10 mouse melanoma cell line was purchased from ATCC (CRL-6475), genetically validated, and maintained in DMEM with 10% (v/v) fetal calf serum and 2 mM glutamine, 100 U/mL penicillin/streptomycin at 37°C, 5% CO2. The cell line was screened for the presence of mycoplasma and mouse pathogens (at Charles River Laboratories, USA) before culturing and never cultured for more than five passages.

### Experimental metastasis assay

B16-F10 (4-5 × 10^5^) cells resuspended in 0.1 mL phosphate buffered saline (PBS) were injected into the tail vein of 6-to 12-week-old sex-matched syngeneic control and mutant mice. After 10 (± 1) days the mice were humanely sacrificed, their lungs removed and washed in PBS and the number of metastatic foci counted macroscopically. The use of the experimental assay as a screen for metastatic colonisation ability has been previously described (Speak et al. 2017).

### Statistics and bioinformatic analysis

The raw data (number of metastatic foci counted in each mutant mouse relative to the wildtype controls) from each cohort of mice was subjected to the non-parametric Mann-Whitney U test. An integrative data analysis (mega-analysis) was performed on the results from all mutant mouse lines that had been tested in ≥ 3 independent cohorts, and was completed using R (package nlme version 3.1) as previously described (van der Weyden et al. 2017b). Using the Mouse Genome Database Informatics (MGI) portal (http://www.informatics.jax.org, v6.14), all 1,300 mutant lines screened were separated into unique symbols and annotated with molecular function using the Gene Ontology (GO) chart tool (Bult et al. 2019) excluding annotations that were Inferred from Electronic Annotation (IEA). Phenotypic information (MP-to-genotype) was pulled from MGI using MouseMine (Motenko et al. 2015) and the phenotypes collapsed to the parental term of the mouse phenotype hierarchy.

### Data availability Statement

Supplemental files available at FigShare. Table S1 lists the targeted genetic regions that were mutated in the genetically modified mice used in the screened. Table S2 is the complete data set (number of metastatic colonies) for all the mice comprising the 1,344 alleles screened (consisting of 23,975 individual mice). Table S3 explains how to interpret the data for the screen supplied in Table S2.

## RESULTS

Tail vein injection of mouse melanoma B16-F10 cells primarily results in pulmonary colonisation (due to the capillary beds in the lungs being the first ones encountered by the cells in the arterial blood after leaving the heart). As these cells are pigmented (melanin granules) their colonisation of the lungs can be determined by macroscopic counting of the number of black metastatic foci on the lungs (**Figure 1A**). Sex-and age-matched wildtype mice were concomitantly dosed with the cohorts of mutant mice (**Figure 1B**), and the results from mutant mice were only compared to the wildtype mice dosed at the same time (due to day-to-day variations in the assay, and factors such as sex and age of the mice affecting metastatic burden (Speak et al. 2017)). A ‘metastatic ratio’ (MR) was determined for each mutant mouse line, which was calculated as the average number of metastatic colonies for the mutant line relative to the average number of metastatic colonies for concomitantly dosed wildtype mice. If a mutant mouse line showed a MR of <0.6 or >1.4 (and Mann-Whitney *P*<0.05), additional cohorts were assayed (n≥2, assayed on independent days). An integrative data analysis (IDA) was performed on the whole dataset and those with *P*<0.005 (Hochberg) and a biological effect (‘genotype effect’) of ≤ −55 or ≥ +55 were classified as ‘hits’. A biological filter was applied as we were only interested in determining robust (strong) regulators of metastatic colonisation.

**Figure 1.**
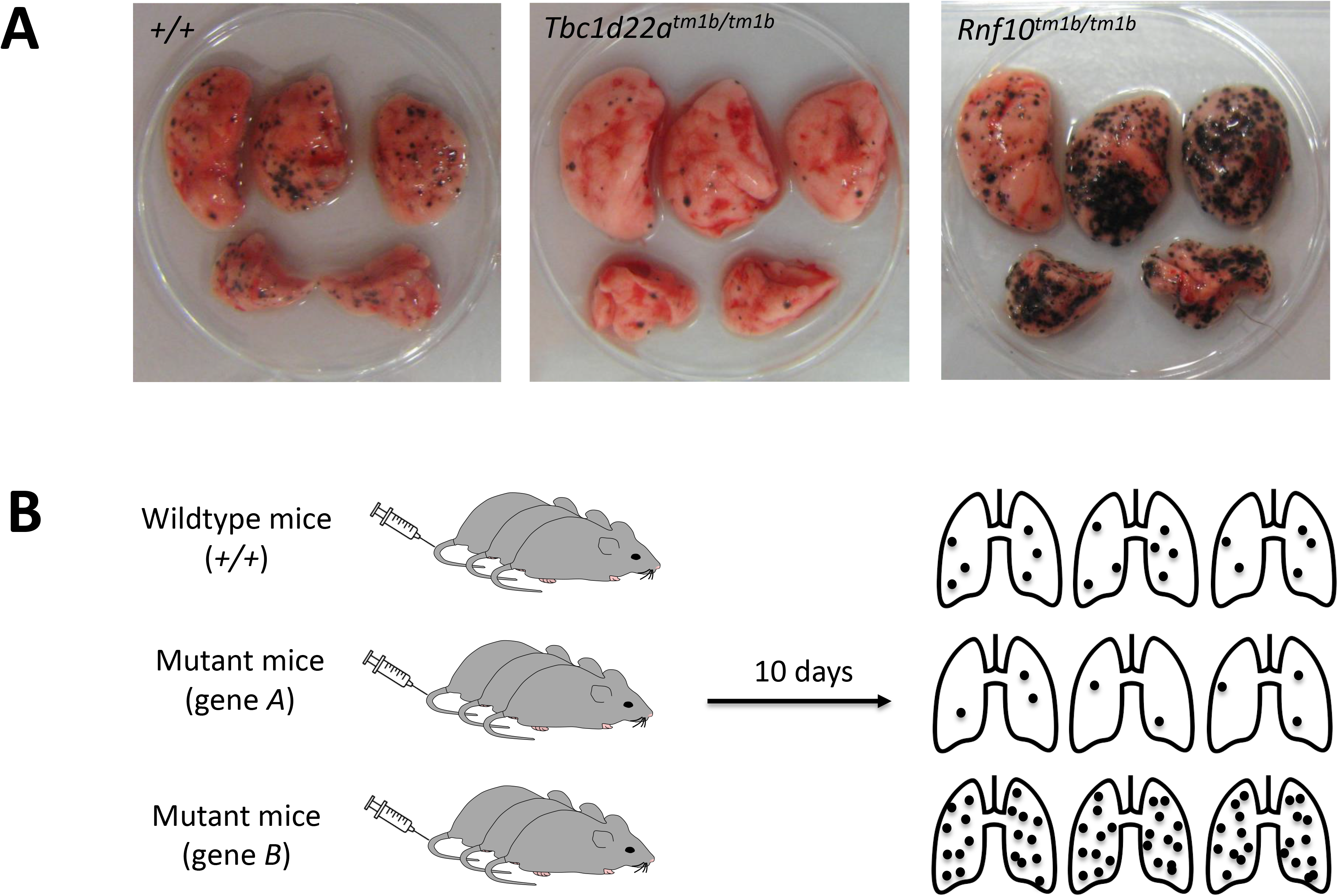
The metastatic colonisation assay. (A) Representative macroscopic image of lungs from wildtype (+/+) and mutant (*Tbc1d22a^tm1b/tm1b^* and *Rnf10^tm1b/tm1b^*) mice 10 days after tail vein dosing with B16-F10 melanoma cells, demonstrating examples of decreased and increased metastatic colonisation, respectively. (B) Schematic of the B16-F10 pulmonary metastasis screen, showing that a cohort of mice consists of wildtype mice and groups of different mutant mice, all of which are tail vein dosed with the B16-F10 melanoma cells, and then the number of pulmonary metastatic colonies counted 10 days later (the ‘metastatic ratio’ of a mutant line is derived by dividing the average of the metastases for a mutant group by the average number of metastases for the wildtype group).

We used *Entpd1* and *Hsp90aa1* mutant mice as positive controls, as the literature suggested they should show altered metastatic burden. *Entpd1* (*ectonucleoside triphosphate diphosphohydrolase 1*) encodes the plasma membrane protein CD39. The enzymatic activity of CD39 (NTPDase I), together with CD73 (ecto-5’-nucleotidase), result in the phosphohydrolysis of extracellular ATP into adenosine, which acts as an immunosuppressive pathway through the activation of adenosine receptors (Stagg and Smyth 2010). *Entpd1*-deficient mice that were administered B16-F10 mouse melanoma cells and MC-38 mouse colon cancer cells via the hepatic portal vein (experimental metastasis assay) were found to develop significantly fewer hepatic metastases than wildtype (control) mice (Sun et al. 2010). In agreement with this, we found that *Entpd1* mutant mice showed significantly reduced numbers of pulmonary metastatic colonies after tail vein dosing with B16-F10 cells, relative to wildtype mice. *Hsp90aa1* (*heat shock protein 90 alpha family class A member 1*) encodes a molecular chaperone that functions to aids in the proper folding of specific target proteins (“clients”), including numerous kinases, transcription factors and steroid hormone receptors (Li et al. 2012). Systemic administration of a mitochondrial-targeted, small-molecule Hsp90 inhibitor (Gamitrinib) to Transgenic Adenocarcinoma of the Mouse Prostate (TRAMP) mice inhibited the formation of localised prostate tumours, as well as the spread of metastatic prostate cancer to abdominal lymph nodes and liver (Kang et al. 2011). In agreement with this, we found that *Hsp90aa1* mutant mice showed significantly reduced numbers of pulmonary metastatic colonies after tail vein dosing with B16-F10 cells, relative to wildtype mice. Both *Entpd1* and *Hsp90aa1* were classified as ‘hits’ using the integrated data analysis (IDA) approach, thus we were confident that our screening methodology was robust.

We have previously published the results of screening 810 mutant mouse lines and showed that endothelial SPNS2 can regulate metastatic colonisation by sphingosine-1-phosphate (S1P)-mediated control of lymphocyte trafficking (van der Weyden et al. 2017a; van der Weyden et al. 2017b). Here we have included an additional 490 mutant mouse lines, to make a total of 1,300 mutant mouse lines (1,344 alleles) screened. The mutant mouse lines were randomly selected and the genes (**Table S1**) cover a diverse range of molecular functions (**Figure 2a**) and are involved in many different biological processes (**Figure 2b**). Of the 1,300 mutant muse lines screened, 1,247 lines (96%) carried alleles that targeted single protein coding genes, with the other alleles targeting lncRNAs (21 lines), miRNAs (8 lines), CpG islands (13 lines), pseudogenes (3 lines), complexes/clusters/regions (3 lines), multiple protein coding genes (3 lines) or gene segments (2 lines). The raw data for each individual mouse (number of metastases counted) is listed in **Table S2**. The mutant mice tested were predominantly homozygotes (880 lines, 68%), with heterozygotes generally only being tested (356 lines, 27%) if the line was lethal or sub-viable (i.e., where 0 or ≤13% of homozygote offspring were obtained from heterozygous intercrosses, respectively) and in a small number of cases both heterozygotes and homozygotes were assessed (64 lines, 5%). An IDA was performed on the data from the 256 mouse lines that showed evidence of a phenotype in initial screening and for which at least 3 independent cohorts were tested, and the 34 mutant lines classified as ‘hits’ are shown **Table 1**.

**Figure 2.**
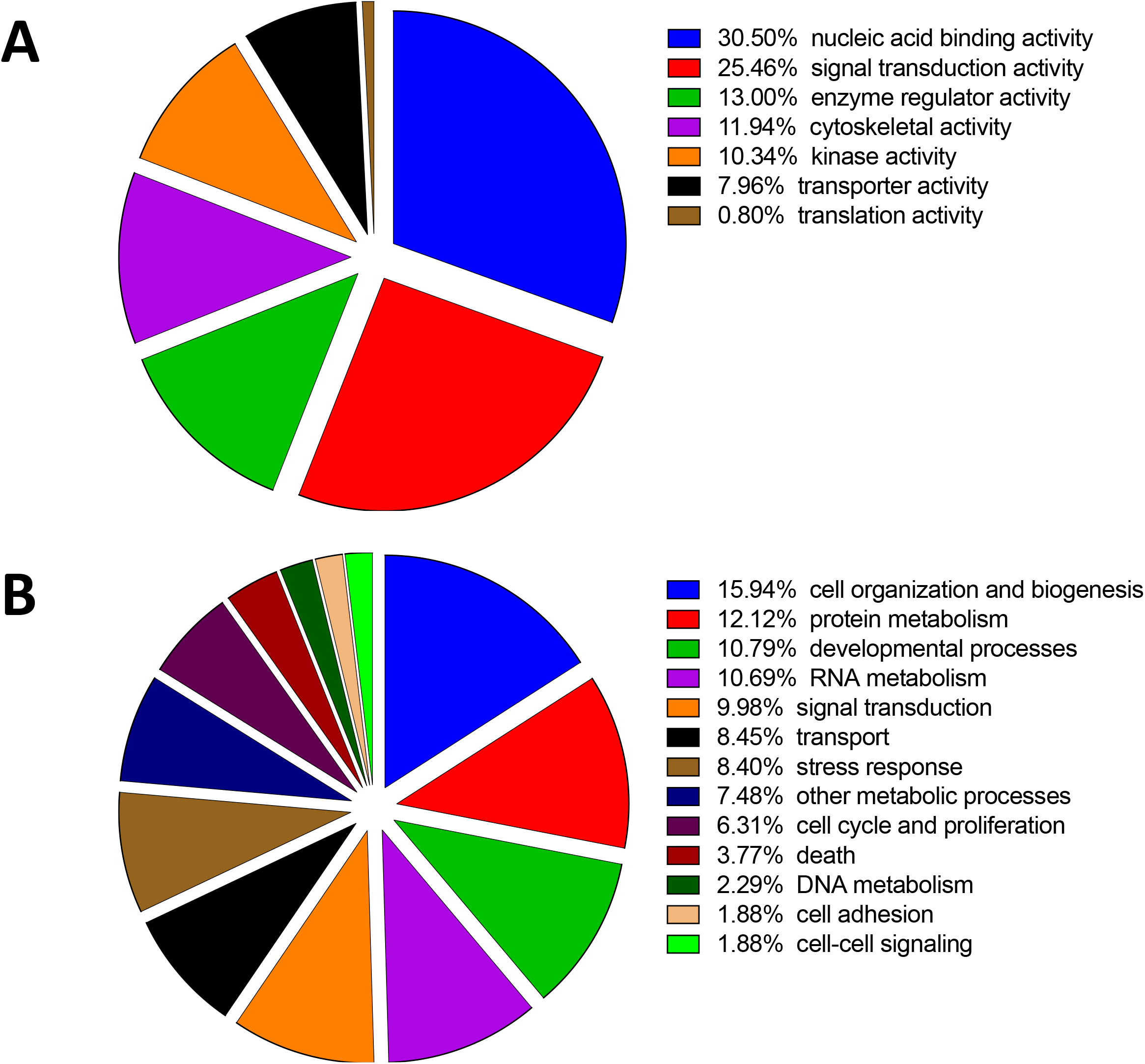
Gene Ontology annotation of the 1,300 mutant mouse lines screened as detailed in Methods. A) Molecular functions of genes screened and B) Biological processes of genes screened.

**Table 1.** Results of the Integrative Data Analysis showing mutant mouse lines with statistically significant decreased or increased pulmonary metastatic colonisation (using a statistical threshold of P<0.0005 and a biological threshold of genotype estimate ≤ −55 or ≥ +55). The genotype of the mutant mice listed were all homozygotes except *Id2*, which were heterozygotes. ‘Cohorts’ was the number of individual cohorts that were tested for this particular mouse line. ‘Genotype estimate’ is the alteration in number of pulmonary metastatic colonies for that mutant mouse line relative to control mice. P value is the Hochberg test. * The mutant mice (and controls) were administered 1/10th dose of other lines. ** The genotype estimate is an average of the values obtained for both sexes (as there was a sex effect observed); the values for each sex are: 290 ± 26 (male) and 153 ± 31 (female).

| GENE | ALLELE | COHORTS | GENOTYPE ESTIMATE | P VALUE |
| --- | --- | --- | --- | --- |
| *Duoxa2* * | <tm1b(KOMP)Wtsi> | 5 | +721 ± 21 | 2.02E-15 |
| *Irf1* | <tm1a(EUCOMM)Wtsi> | 4 | +246 ± 32 | 1.12E-04 |
| *Rnf10* ** | <tm1b(KOMP)Wtsi> | 5 | +221 ± 28 | 1.74E-08 |
| *Pik3cg* | <tm1a(EUCOMM)Wtsi> | 6 | +198 ± 14 | 7.20E-12 |
| *Herc1* | <em1Wtsi> | 4 | +178 ± 27 | 2.75E-04 |
| *Arhgap30* | <tm1a(EUCOMM)Wtsi> | 5 | +178 ± 11 | 2.25E-31 |
| *Dph6* | <tm1a(KOMP)Wtsi> | 5 | +141 ± 13 | 7.94E-17 |
| *Slc9a3r2* | <tm1a(EUCOMM)Wtsi> | 6 | +141 ± 20 | 2.11E-05 |
| *Igh-6* | <tm1(Cgn)> | 6 | +101 ± 13 | 6.78E-08 |
| *Abhd17a* | <tm1a(KOMP)Wtsi> | 6 | +96 ± 11 | 1.99E-08 |
| *Irf7* | <tm1(KOMP)Wtsi> | 6 | +92 ± 16 | 5.39E-05 |
| *Id2* | <tm2b(EUCOMM)Wtsi> | 6 | +89 ± 11 | 4.91E-12 |
| *Fgd4* | <tm1a(EUCOMM)Wtsi> | 4 | +89 ± 15 | 1.15E-03 |
| *Ankhd1* | <tm1b(KOMP)Wtsi> | 5 | +60 ± 10 | 1.68E-06 |
| *Fzd6* | <tm2a(EUCOMM)Wtsi> | 6 | +59 ± 10 | 7.70E-07 |
| *Grsf1* | <tm1b(EUCOMM)Wtsi> | 7 | -55 ± 5 | 6.36E-12 |
| *Ncf4* | <tm2Pth> | 8 | -61 ± 9 | 3.84E-08 |
| *Mier1* | <tm1a(EUCOMM)Wtsi> | 3 | -63 ± 13 | 5.02E-04 |
| *Cybc1* | <tm1a(KOMP)Wtsi> | 11 | -67 ± 11 | 1.79E-06 |
| *Abraxas2* | <tm1a(EUCOMM)Wtsi> | 3 | -67 ± 11 | 4.42E-04 |
| *Rwdd1* | <tm1b(KOMP)Wtsi> | 9 | -70 ± 8 | 1.79E-14 |
| *Cybb* | <tm1Din> | 6 | -71 ± 8 | 2.76E-12 |
| *Bach2* | <tm1a(EUCOMM)Wtsi> | 6 | -71 ± 7 | 7.12E-10 |
| *Ncf1* | <m1J> | 3 | -74 ± 10 | 8.51E-06 |
| *Ncf2* | <tm1a(EUCOMM)Wtsi> | 6 | -77 ± 6 | 5.99E-13 |
| *Lrig1* | <tm1a(EUCOMM)Wtsi> | 5 | -92 ± 10 | 2.53E-13 |
| *Cyba* | <tm1a(EUCOMM)Wtsi> | 6 | -99 ± 6 | 1.18E-17 |
| *Arhgef1* | <tm1a(EUCOMM)Wtsi> | 6 | -100 ± 6 | 7.03E-14 |
| *Fbxo7* | <tm1a(EUCOMM)Wtsi> | 4 | -100 ± 15 | 7.63E-08 |
| *Tbc1d22a* | <tm1b(KOMP)Wtsi> | 6 | -102 ± 9 | 5.12E-22 |
| *Hsp90aa1* | <tm1(KOMP)Wtsi> | 3 | -103 ± 13 | 6.37E-06 |
| *Entpd1* | <tm1a(EUCOMM)Wtsi/Hmgu> | 4 | -106 ± 16 | 9.13E-08 |
| *Nbeal2* | <tm1a(EUCOMM)Wtsi> | 4 | -124 ± 14 | 9.79E-09 |
| *Spns2* | <tm1a(KOMP)Wtsi> | 10 | -160 ± 6 | 2.81E-37 |

## DISCUSSION

We have characterised the mechanism of action for several genes that showed a decreased metastatic colonisation phenotype in our screen, specifically *Spns2* (van der Weyden et al. 2017a), *Nbeal2* (Guerrero et al. 2014), *Cybc1* (Thomas et al. 2017) and the 5 members of the NOX2 complex (van der Weyden et al. 2018). These genes regulate pulmonary metastatic colonisation primarily by impacting on the function of the haematopoietic/immune system (lymphocytes, granulocytes/monocytes and platelets). In addition, *Bach2* was also a ‘hit’ in our screen, showing decreased metastatic colonisation, and Bach2 is a key regulator of CD4^+^ T-cell differentiation (Roychoudhuri et al. 2013), with *Bach2* mutant mice recently being shown to have increased CD8^+^ T-cell cytotoxic activity (Abeler-Dorner et al. 2020). Indeed, phenotypic analysis of the 34 metastatic colonisation regulating genes detailed in this study show a strong enrichment for genes involved in immune/haematopoietic system development and function with phenotypes in those categories representing 88% of all the reported phenotypes associated with those genes.

As we have previously predominantly focussed our attention genes positively regulating metastatic colonisation (i.e., mutant mice showing decreased metastasis), we will now turn our focus to discussing negative regulators of metastatic colonisation (i.e., mutant mice showing increased metastasis).

### Duoxa2

The strongest biological phenotype we observed, in terms of increased numbers of pulmonary metastatic colonies relative to controls, was with *Duoxa2* mutant mice. The *Dual oxidase maturation factor 2 (DUOXA2*) gene encodes an endoplasmic reticulum protein that is necessary for the maturation and cellular localization (transport from the endoplasmic reticulum to the plasma membrane) of dual oxidase 2 (DUOX2) (Grasberger and Refetoff 2006). The NADPH oxidases, DUOX1 and DUOX2, are critical for the production of extracellular hydrogen peroxide that is required for thyroperoxidase-mediated thyroid hormone synthesis in the thyroid gland; as a result, mutations in DUOX and/or DUOXA2 result in thyroid dyshormonogenesis and congenital hypothyroidism (De Deken and Miot 2019). Indeed, *Duoxa2* mutant mice were significantly smaller than their wildtype or heterozygous littermates. A smaller body size (and thus total blood volume) undoubtedly contributed to the increased pulmonary metastatic burden we observed, as well as the presence of extrapulmonary metastases (bone marrow, liver, kidney). However, it was recently shown that *Duoxa2* mutant mice have alterations in key immune cell subsets (CD4+ T-cells, neutrophils, monocytes and NK cells) (Abeler-Dorner et al. 2020), which could also account for their increased metastatic colonisation. Thus, generation of an inducible *Duoxa2* mutant mouse, wherein *Duoxa2* could be deleted in an adult mouse, will be required to disentangle any effects that loss of DUOXA2 may be having on metastatic colonisation aside from a smaller body size.

### Rnf10

The *Ring finger protein 10* (*Rnf10*) gene encodes a protein with a ring finger motif (a C3HC4-type zinc finger). *Rnf10*/RNF10 has been shown to be important for key neurobiology functions, including myelin formation (Hoshikawa et al. 2008), neuronal cell differentiation (Malik et al. 2013) and synaptonuclear messaging (Carrano et al. 2019). It has also been reported to play a role in vascular restenosis (Li et al. 2019) and a SNP in *RNF10* has been associated with adiposity and type 2 diabetes (Huang et al. 2014). We found that *Rnf10-*deficient mice showed increased pulmonary metastatic colonisation, with males having a consistently higher metastatic burden than females (290 ± 26 versus 153 ± 31, respectively); this is the only mutant line in which we observed a sexually dimorphic effect. Further investigations are required to provide mechanistic insight as to how *Rnf10* may be regulating metastatic colonisation, and why it has a stronger effect in males.

### Slc9a3r2

The *Slc9a3r2* (*SLC9A3 regulator 2*) gene encodes a member of the Na(+)/H(+) exchanger regulatory factor (NHERF) family of PDZ scaffolding proteins. All NHERF proteins are involved with anchoring membrane proteins that contain PDZ recognition motifs to form multiprotein signalling complexes. SLC9A3R2 (also known as NHERF2) has been shown to form complexes with a diverse range of proteins depending on tissue context, including complexing with the lysophosphatidic acid (LPA) receptor and the epithelial anion channel, cystic fibrosis transmembrane conductance regulator (CFTR) in airway and gut epithelia (Zhang et al. 2017), the P2Y1 nucleotide and mGluR5 glutamate receptors to different ion channels in neurons (Filippov et al. 2010) and megalin and ClC-5 in proximal tubule cells (Hryciw et al. 2012). However, it is not yet clear how loss of *Slc9a3r* results in increased metastatic colonisation.

### *Ankhd1* and *Fzd6*

The *Ankhd1* (ANKHD1 ankyrin repeat and KH domain containing 1) gene encodes a protein with multiple ankyrin repeat domains and a single KH-domain. ANKHD1 is the mammalian homolog of *Mask1* in Drosophila (which is required for the activity of the Hippo pathway effector, Yorkie) and promotes YAP1 activation and cell cycle progression (Machado-Neto et al. 2014). Studies have demonstrated a role for ANKHD1 in promoting cell cycle progression/proliferation in renal cancer, multiple myeloma cell and prostate cancer cells (Dhyani et al. 2012; Machado-Neto et al. 2014; Fragiadaki and Zeidler 2018) and in promoting hepatocellular carcinoma metastasis (Zhou et al. 2019). The *Fzd6* (frizzled class receptor 6) gene is a member of the ‘frizzled’ gene family, which encode 7-transmembrane domain proteins that are receptors for Wnt signaling proteins. FDZ6 has a known role in non-canonical WNT/PCP signalling in cancer (Corda and Sala 2017), including mediating transformation, increased invasiveness of tumour cells and metastasis (Cantilena et al. 2011; Corda and Sala 2017; Corda et al. 2017). In agreement with this, increased expression of FDZ6 has reported in many cancer types, and correlates with poor prognosis in patients with breast, brain and oesophageal cancer (Corda et al. 2017; Huang et al. 2016; Zhang et al. 2019). Thus, both *Ankhd1* and Fzd6 have well-characterised tumour cell-intrinsic roles, however, how they mediate their role in regulating tumour cell extrinsic metastasis is not clear.

### Regulation of the immune system

Five genes with known roles in regulating immune cell function were identified; the immune cell types included NK cells, lymphocytes (T-and B-cells) and granulocytes/macrophages, which have all been shown to play critical roles in regulating metastasis (reviewed in (Blomberg et al. 2018)). Thus genes that interfere with their production, maturation and/or function could understandably result in increased levels of metastatic colonisation.

#### Irf1

The *Interferon regulatory factor 1* (*IRF1*) gene encodes a transcription factor that is one of 9 members of the interferon regulatory transcription factor (IRF) family. IRF1 stimulates both innate and acquired immune responses through the activation of specific target genes, by regulating target genes through binding to an interferon-stimulated response element (ISRE) in their promoters and inducing either transcriptional activation or repression (Ikushima et al. 2013). *Irf1* null mice are immunodeficient, characterised by a marked reduction in CD8^+^ T cells (Penninger et al. 1997) and a decrease in NK cell numbers with associated impaired cytolytic activity (Taki et al. 1997).

#### Irf7

The *Interferon regulatory factor 7* (*IRF7*) gene is another member of the IRF family and plays a critical role in the innate immune response against viruses. *Irf7-*null mice are highly susceptible to H1N1 infection (Wilk et al. 2015) and secrete decreased levels of IFN-α/β in response to stimulation (Honda et al. 2005).

#### Id2

The *ID2* (*inhibitor of DNA binding 2*) gene encodes a helix-loop-helix-containing protein that lacks a DNA-binding domain and is one of the four members of the ID family (ID1– ID4). ID proteins dimerise with E protein, RB and Ets transcription factors, preventing the formation of DNA-binding transcription complexes. *Id2* null mice show a greatly reduced population of natural killer (NK) cells, as *Id2* plays a role in NK cell maturation (Yokota et al. 1999; Boos et al. 2007).

#### Igh-6

The *IGH-6* (*immunoglobulin heavy constant mu*) gene encodes a protein that is important for the production of the heavy chain of IgM antibodies and maturation of pre-B cells, the precursors of B-lymphocytes. *Igh-6* null mice are B-cell-deficient, with their development arrested at the stage of pre-B-cell maturation (Kitamura et al. 1991). *Igh-6*^-/-^ mice also show impaired Th1 T-cell responses to Salmonella antigens/infections (Mastroeni et al. 2000; Ugrinovic et al. 2003) demonstrating a role for B cells in the establishment and/or persistence of a stable T-cell memory pool.

#### Pik3cg

The *PIK3CG* (*phosphatidylinositol-4,5-bisphosphate 3-kinase catalytic subunit gamma*) gene, encodes a class I catalytic subunit of phosphoinositide 3-kinase (PI3K), known by many names, including p110-γ and PI3Kγ. Like other class I catalytic subunits (p110-*α*, p110-*β*, and p110-δ), p110-γ binds a p85 regulatory subunit to form PI3K, which phosphorylate inositol lipids and is involved in the immune response. p110-γ is highly expressed in leukocytes and is important for restraining inflammation and promoting appropriate adaptive immune responses in both humans and mice (Takeda et al. 2019). *p110-γ* null mice show defective thymocyte development and T cell activation, as well as neutrophil migration and oxidative burst (Sasaki et al. 2000).

### Protein modification

A number of genes encoding protein modifiers were identified; these included a ubiquitin-related protein, a serine hydrolase and a protein involved in amidation. How the targets of these proteins can regulate metastatic colonisation is still unclear.

#### Herc1

The *HERC1* (*HECT and RLD domain containing E3 ubiquitin protein ligase family member 1*) gene encodes an E3 ubquitin ligase protein. In humans, six *HERC* genes have been reported which encode two subgroups of HERC proteins: large (HERC1-2) and small (HERC3-6). The HERC1 protein was the first to be identified and has been found to play numerous roles, including membrane trafficking, protein stability and DNA damage repair, through its interactions with clathrin, TSC2 and pyruvate kinase (M2 isoform), respectively (reviewed in (Garcia-Cano et al. 2019)). *Tambaleante* (*tbl*) mice, which carry a spontaneous missense mutation in *Herc1*, show neurological phenotypes including, Purkinje cell degeneration, hind limb clasping and impaired rotarod performance (Mashimo et al. 2009). Whilst mutations/loss of *HERC1* expression have been reported in some cancers (reviewed in (Garcia-Cano et al. 2019)), it is not clear how tumour cell-extrinsic loss of *Herc1* resulted in increased metastatic colonisation.

#### Abhd17a

The *ABHD17A* (*abhydrolase domain containing 17A*) gene encodes a member of the ABHD17 family of proteins that are membrane-anchored serine hydrolases which can accelerate palmitate turnover on PSD-95 and N-Ras. The catalytic activity of ABHD17 proteins are required for N-Ras depalmitoylation and re-localization to internal cellular membranes (Lin and Conibear 2015) and ABHD17 proteins finely control the amount of synaptic PSD-95 by regulating PSD-95 palmitoylation cycles in neurons (Yokoi et al. 2016). More recently, regulation of the palmitoylation status of the transcription factor TEAD, which is depalmitoylased by ABHD17A, has been suggested to be a potential target for controlling TEAD-dependent processes, including cancer cell growth (Kim and Gumbiner 2019).

#### Dph6

The *Dph6* (*diphthamine biosynthesis 6*) gene encodes protein that is required for the amidation step of the diphthamide pathway in yeast. Diphthamide is a highly modified histidine residue in eukaryotic translation elongation factor 2 (eEF2) and diphthamide synthesis is required for optimal translational accuracy and cell growth (Uthman et al. 2013). In eukaryotes, the formation of diphthamide involves a conserved biosynthetic pathway involving 7 members, DPH1-7 that has been predominantly studied in yeast (reviewed in (Schaffrath et al. 2014)). However, they do play an import role in mammalian cells as *Dph1* null mice display multiple developmental defects that parallel Miller-Dieker syndrome (MDS) (Yu et al. 2014), associated with deletions on chromosome 17p13.3, *Dph3* null mice are embryonically lethal (Liu et al. 2006) and *Dnajc24* (*Dph4*) null mice almost always die before birth with the few that do survive showing severe developmental defects reminiscent of *Dph1* null mice (Webb et al. 2008). Recently, *Dph6* mutant mice were shown to have an immune phenotype with alterations in many innate and adaptive cell lineages (Abeler-Dorner et al. 2020), and it is possible that these may be affecting metastatic colonisation. Thus, as very little is known about ABHD17A and DPH6 in the context of cancer, it is difficult to precisely speculate how they may be playing a role in tumour cell extrinsic regulation of metastatic colonisation.

### Rho GTPase regulating proteins

Rho GTPases are molecular switches that control a wide variety of signal-transduction pathways, including regulation of the cytoskeleton, migration, and proliferation. Rho GTPases can be regulated by GTPase-activating proteins (GAPs) and Rho GDP/GTP nucleotide exchange factors (Rho GEFs). We identified one of each of these family members.

#### Arhgap30

The *ARHGAP30* (*Rho GTPase activating protein 30*) gene encodes a Rho GTPase-activating protein, with a role in regulating cell adhesion (Naji et al. 2011), as well as suppressing lung cancer cell proliferation, migration and invasion (Mao and Tong 2018). In colorectal cancer (CRC), ARHGAP30 levels correlate with p53 acetylation and functional activation (Wang et al. 2014), and ARHGAP30 has been proposed as a prognostic marker for CRC (Wang et al. 2014), early-stage pancreatic ductal adenocarcinoma (Liao et al. 2017) and lung adenocarcinoma (Li et al. 2018).

#### Fgd4

The *FDG4* (*FYVE, RhoGEF And PH Domain Containing 4*) gene encodes a GEF specific to the Rho GTPase, CDC42. FDG4, also known as FRABIN, contains an actin filament-binding domain (ABD), an Dbl homology domain (DHD), a cysteine rich-domain (CRD), and two pleckstrin homology domains (PHD), which are involved in binding to the actin and activating CDC42 at that vicinity, resulting in actin cytoskeleton reorganization (allowing for shape changes such as the formation of filopodia and lamellipodia) (Nakanishi and Takai 2008). FGD4 overexpression has been observed in pancreatic neuroendocrine neoplasms (Shahid et al. 2019) and expression of FDG4 positively correlates with the aggressive phenotype of prostate cancer (Bossan et al. 2018). Mutations in this gene can cause Charcot-Marie-Tooth (CMT) disease type 4H (CMT4H), characterised by heterogeneous hereditary motor and sensory neuropathies as a result of demyelination of peripheral nerves (Delague et al. 2007).

Although much is known about these two genes and they have well established roles in tumour cell-intrinsic roles in cancer, it is not clear at this stage how they may be mediating an increased metastatic colonisation phenotype.

## CONCLUSION

In summary, we have used the experimental metastasis assay to screen 1,300 mutant mouse lines to identify novel host/microenvironmental regulators of metastatic colonisation. We have identified 34 genes whose loss of expression results in either an increased or decreased ability for mouse melanoma cells to undergo metastatic colonisation of the lung following tail vein injection. Some of these genes regulate key pathways in immune cell development or function, however many have only been shown to play a role in tumour cell-intrinsic pathways with no known tumour cell-extrinsic functions reported, thus, we have identified numerous novel regulators of pulmonary metastatic colonisation, which could represent potential therapeutic targets.

## ACKNOWLEDGEMENTS

We thank the Mouse pipeline teams and the Research Support Facility staff at the Wellcome Sanger Institute for the generation, genotyping and care of the mice used in this study. This work was funded by the Wellcome Trust (awarded to D.J.A.).

## AUTHOR CONTRIBUTIONS

L.v.d.W, A.O.S and D.J.A conceived the experiments and led the work. L.v.d.W, A.S., V.I. and A.O.S performed the experiments and interpreted the data. L.v.d.W. wrote the manuscript and all authors critically reviewed the manuscript and approved the final version.

## SUPPLEMENTARY DATA

**Table S1. A list of the targeted genetic regions that were mutated in the genetically modified mice used in the screened.** The name, feature type and chromosomal location is listed for all the genes/ genetic locations that were targeted in the mutant mice screened. ‘Feature type’ is as defined at http://www.informatics.jax.org/userhelp/GENE_feature_types_help.shtml. The ‘current symbol’ is the most recent name for the gene/genetic region (as on some occasions the name has been changed at MGI since the mouse line was originally screened).

**Table S2. Complete data set for all the mice comprising the 1,344 alleles screened.** Results of the B16-F10 pulmonary metastasis screen for the 1,344 alleles consisting of 23,975 individual mice. There are a number of meta-data (such as sex, genetic background, assay date, zygosity, gene, cell number used and QC notes) that should be considered in any analysis. Details of how to interpret the data are explained in Table S3.

**Table S3. Explanation of how to interpret the data for the screen.** An explanation of the data format, data options and other relevant information to allow interpretation of the data in Table S2.

## REFERENCES

Abeler-Dorner, L., A.G. Laing, A. Lorenc, D.S. Ushakov, S. Clare et al., 2020 High-throughput phenotyping reveals expansive genetic and structural underpinnings of immune variation. Nat Immunol 21 (1):86–100.

Blomberg, O.S., L. Spagnuolo, and K.E. de Visser, 2018 Immune regulation of metastasis: mechanistic insights and therapeutic opportunities. Dis Model Mech 11 (10).

Boos, M.D., Y. Yokota, G. Eberl, and B.L. Kee, 2007 Mature natural killer cell and lymphoid tissue-inducing cell development requires Id2-mediated suppression of E protein activity. J Exp Med 204 (5):1119–1130.

Bossan, A., R. Ottman, T. Andl, M.F. Hasan, N. Mahajan et al., 2018 Expression of FGD4 positively correlates with the aggressive phenotype of prostate cancer. BMC Cancer 18 (1):1257.

Bult, C.J., J.A. Blake, C.L. Smith, J.A. Kadin, J.E. Richardson et al., 2019 Mouse Genome Database (MGD) 2019. Nucleic Acids Res 47 (D1):D801–D806.

Cantilena, S., F. Pastorino, A. Pezzolo, O. Chayka, V. Pistoia et al., 2011 Frizzled receptor 6 marks rare, highly tumourigenic stem-like cells in mouse and human neuroblastomas. Oncotarget 2 (12):976–983.

Carrano, N., T. Samaddar, E. Brunialti, L. Franchini, E. Marcello et al., 2019 The Synaptonuclear Messenger RNF10 Acts as an Architect of Neuronal Morphology. Mol Neurobiol 56 (11):7583–7593.

Chambers, A.F., G.N. Naumov, H.J. Varghese, K.V. Nadkarni, I.C. MacDonald et al., 2001 Critical steps in hematogenous metastasis: an overview. Surg Oncol Clin N Am 10 (2):243–255, vii.

Corda, G., and A. Sala, 2017 Non-canonical WNT/PCP signalling in cancer: Fzd6 takes centre stage. Oncogenesis 6 (7):e364.

Corda, G., G. Sala, R. Lattanzio, M. Iezzi, M. Sallese et al., 2017 Functional and prognostic significance of the genomic amplification of frizzled 6 (FZD6) in breast cancer. J Pathol 241 (3):350–361.

De Deken, X., and F. Miot, 2019 DUOX Defects and Their Roles in Congenital Hypothyroidism. Methods Mol Biol 1982:667–693.

Delague, V., A. Jacquier, T. Hamadouche, Y. Poitelon, C. Baudot et al., 2007 Mutations in FGD4 encoding the Rho GDP/GTP exchange factor FRABIN cause autosomal recessive Charcot-Marie-Tooth type 4H. Am J Hum Genet 81 (1):1–16.

Dhyani, A., A.S. Duarte, J.A. Machado-Neto, P. Favaro, M.M. Ortega et al., 2012 ANKHD1 regulates cell cycle progression and proliferation in multiple myeloma cells. FEBS Lett 586 (24):4311–4318.

Fidler, I.J., and M.L. Kripke, 2015 The challenge of targeting metastasis. Cancer Metastasis Rev 34 (4):635–641.

Filippov, A.K., J. Simon, E.A. Barnard, and D.A. Brown, 2010 The scaffold protein NHERF2 determines the coupling of P2Y1 nucleotide and mGluR5 glutamate receptor to different ion channels in neurons. J Neurosci 30 (33):11068–11072.

Fragiadaki, M., and M.P. Zeidler, 2018 Ankyrin repeat and single KH domain 1 (ANKHD1) drives renal cancer cell proliferation via binding to and altering a subset of miRNAs. J Biol Chem 293 (25):9570–9579.

Garcia-Cano, J., A. Martinez-Martinez, J. Sala-Gaston, L. Pedrazza, and J.L. Rosa, 2019 HERCing: Structural and Functional Relevance of the Large HERC Ubiquitin Ligases. Front Physiol 10:1014.

Grasberger, H., and S. Refetoff, 2006 Identification of the maturation factor for dual oxidase. Evolution of an eukaryotic operon equivalent. J Biol Chem 281 (27):18269–18272.

Guerrero, J.A., C. Bennett, L. van der Weyden, H. McKinney, M. Chin et al., 2014 Gray platelet syndrome: proinflammatory megakaryocytes and alpha-granule loss cause myelofibrosis and confer metastasis resistance in mice. Blood 124 (24):3624–3635.

Honda, K., H. Yanai, H. Negishi, M. Asagiri, M. Sato et al., 2005 IRF-7 is the master regulator of type-I interferon-dependent immune responses. Nature 434 (7034):772–777.

Hoshikawa, S., T. Ogata, S. Fujiwara, K. Nakamura, and S. Tanaka, 2008 A novel function of RING finger protein 10 in transcriptional regulation of the myelin-associated glycoprotein gene and myelin formation in Schwann cells. PLoS One 3 (10):e3464.

Hryciw, D.H., K.A. Jenkin, A.C. Simcocks, E. Grinfeld, A.J. McAinch et al., 2012 The interaction between megalin and ClC-5 is scaffolded by the Na(+)-H(+) exchanger regulatory factor 2 (NHERF2) in proximal tubule cells. Int J Biochem Cell Biol 44 (5):815–823.

Huang, K., A.K. Nair, Y.L. Muller, P. Piaggi, L. Bian et al., 2014 Whole exome sequencing identifies variation in CYB5A and RNF10 associated with adiposity and type 2 diabetes. Obesity (Silver Spring) 22 (4):984–988.

Huang, T., A.A. Alvarez, R.P. Pangeni, C.M. Horbinski, S. Lu et al., 2016 A regulatory circuit of miR-125b/miR-20b and Wnt signalling controls glioblastoma phenotypes through FZD6-modulated pathways. Nat Commun 7:12885.

Ikushima, H., H. Negishi, and T. Taniguchi, 2013 The IRF family transcription factors at the interface of innate and adaptive immune responses. Cold Spring Harb Symp Quant Biol 78:105–116.

Kang, B.H., M. Tavecchio, H.L. Goel, C.C. Hsieh, D.S. Garlick et al., 2011 Targeted inhibition of mitochondrial Hsp90 suppresses localised and metastatic prostate cancer growth in a genetic mouse model of disease. Br J Cancer 104 (4):629–634.

Kim, N.G., and B.M. Gumbiner, 2019 Cell contact and Nf2/Merlin-dependent regulation of TEAD palmitoylation and activity. Proc Natl Acad Sci U S A 116 (20):9877–9882.

Kitamura, D., J. Roes, R. Kuhn, and K. Rajewsky, 1991 A B cell-deficient mouse by targeted disruption of the membrane exon of the immunoglobulin mu chain gene. Nature 350 (6317):423–426.

Leach, D.R., M.F. Krummel, and J.P. Allison, 1996 Enhancement of antitumor immunity by CTLA-4 blockade. Science 271 (5256):1734–1736.

Li, J., J. Soroka, and J. Buchner, 2012 The Hsp90 chaperone machinery: conformational dynamics and regulation by co-chaperones. Biochim Biophys Acta 1823 (3):624–635.

Li, L., M. Peng, W. Xue, Z. Fan, T. Wang et al., 2018 Integrated analysis of dysregulated long non-coding RNAs/microRNAs/mRNAs in metastasis of lung adenocarcinoma. J Transl Med 16 (1):372.

Li, S., G. Yu, F. Jing, H. Chen, A. Liu et al., 2019 RING finger protein 10 attenuates vascular restenosis by inhibiting vascular smooth muscle cell hyperproliferation in vivo and vitro. IUBMB Life 71 (5):632–642.

Liao, X., K. Huang, R. Huang, X. Liu, C. Han et al., 2017 Genome-scale analysis to identify prognostic markers in patients with early-stage pancreatic ductal adenocarcinoma after pancreaticoduodenectomy. Onco Targets Ther 10:4493–4506.

Lin, D.T., and E. Conibear, 2015 ABHD17 proteins are novel protein depalmitoylases that regulate N-Ras palmitate turnover and subcellular localization. Elife 4:e11306.

Liu, S., J.F. Wiggins, T. Sreenath, A.B. Kulkarni, J.M. Ward et al., 2006 Dph3, a small protein required for diphthamide biosynthesis, is essential in mouse development. Mol Cell Biol 26 (10):3835–3841.

Machado-Neto, J.A., M. Lazarini, P. Favaro, G.C. Franchi, Jr., A.E. Nowill et al., 2014 ANKHD1, a novel component of the Hippo signaling pathway, promotes YAP1 activation and cell cycle progression in prostate cancer cells. Exp Cell Res 324 (2):137–145.

Malik, Y.S., M.A. Sheikh, M. Lai, R. Cao, and X. Zhu, 2013 RING finger protein 10 regulates retinoic acid-induced neuronal differentiation and the cell cycle exit of P19 embryonic carcinoma cells. J Cell Biochem 114 (9):2007–2015.

Mao, X., and J. Tong, 2018 ARHGAP30 suppressed lung cancer cell proliferation, migration, and invasion through inhibition of the Wnt/beta-catenin signaling pathway. Onco Targets Ther 11:7447–7457.

Mashimo, T., O. Hadjebi, F. Amair-Pinedo, T. Tsurumi, F. Langa et al., 2009 Progressive Purkinje cell degeneration in tambaleante mutant mice is a consequence of a missense mutation in HERC1 E3 ubiquitin ligase. PLoS Genet 5 (12):e1000784.

Mastroeni, P., C. Simmons, R. Fowler, C.E. Hormaeche, and G. Dougan, 2000 Igh-6(-/-) (B-cell-deficient) mice fail to mount solid acquired resistance to oral challenge with virulent Salmonella enterica serovar typhimurium and show impaired Th1 T-cell responses to Salmonella antigens. Infect Immun 68 (1):46–53.

Meehan, T.F., N. Conte, D.B. West, J.O. Jacobsen, J. Mason et al., 2017 Disease model discovery from 3,328 gene knockouts by The International Mouse Phenotyping Consortium. Nat Genet 49 (8):1231–1238.

Motenko, H., S.B. Neuhauser, M. O’Keefe, and J.E. Richardson, 2015 MouseMine: a new data warehouse for MGI. Mamm Genome 26 (7-8):325–330.

Naji, L., D. Pacholsky, and P. Aspenstrom, 2011 ARHGAP30 is a Wrch-1-interacting protein involved in actin dynamics and cell adhesion. Biochem Biophys Res Commun 409 (1):96–102.

Nakanishi, H., and Y. Takai, 2008 Frabin and other related Cdc42-specific guanine nucleotide exchange factors couple the actin cytoskeleton with the plasma membrane. J Cell Mol Med 12 (4):1169–1176.

Penninger, J.M., C. Sirard, H.W. Mittrucker, A. Chidgey, I. Kozieradzki et al., 1997 The interferon regulatory transcription factor IRF-1 controls positive and negative selection of CD8+ thymocytes. Immunity 7 (2):243–254.

Quail, D.F., and J.A. Joyce, 2013 Microenvironmental regulation of tumor progression and metastasis. Nat Med 19 (11):1423–1437.

Roychoudhuri, R., K. Hirahara, K. Mousavi, D. Clever, C.A. Klebanoff et al., 2013 BACH2 represses effector programs to stabilize T(reg)-mediated immune homeostasis. Nature 498 (7455):506–510.

Sasaki, T., J. Irie-Sasaki, R.G. Jones, A.J. Oliveira-dos-Santos, W.L. Stanford et al., 2000 Function of PI3Kgamma in thymocyte development, T cell activation, and neutrophil migration. Science 287 (5455):1040–1046.

Schaffrath, R., W. Abdel-Fattah, R. Klassen, and M.J. Stark, 2014 The diphthamide modification pathway from Saccharomyces cerevisiae--revisited. Mol Microbiol 94 (6):1213–1226.

Shahid, M., T.B. George, J. Saller, M. Haija, Z. Sayegh et al., 2019 FGD4 (Frabin) Overexpression in Pancreatic Neuroendocrine Neoplasms. Pancreas 48 (10):1307–1311.

Speak, A.O., A. Swiatkowska, N.A. Karp, M.J. Arends, D.J. Adams et al., 2017 A high-throughput in vivo screening method in the mouse for identifying regulators of metastatic colonization. Nat Protoc 12 (12):2465–2477.

Stagg, J., and M.J. Smyth, 2010 Extracellular adenosine triphosphate and adenosine in cancer. Oncogene 29 (39):5346–5358.

Sun, X., Y. Wu, W. Gao, K. Enjyoji, E. Csizmadia et al., 2010 CD39/ENTPD1 expression by CD4+Foxp3+ regulatory T cells promotes hepatic metastatic tumor growth in mice. Gastroenterology 139 (3):1030–1040.

Takeda, A.J., T.J. Maher, Y. Zhang, S.M. Lanahan, M.L. Bucklin et al., 2019 Human PI3Kgamma deficiency and its microbiota-dependent mouse model reveal immunodeficiency and tissue immunopathology. Nat Commun 10 (1):4364.

Taki, S., T. Sato, K. Ogasawara, T. Fukuda, M. Sato et al., 1997 Multistage regulation of Th1-type immune responses by the transcription factor IRF-1. Immunity 6 (6):673–679.

Talmadge, J.E., and I.J. Fidler, 2010 AACR centennial series: the biology of cancer metastasis: historical perspective. Cancer Res 70 (14):5649–5669.

Thomas, D.C., S. Clare, J.M. Sowerby, M. Pardo, J.K. Juss et al., 2017 Eros is a novel transmembrane protein that controls the phagocyte respiratory burst and is essential for innate immunity. J Exp Med 214 (4):1111–1128.

Ugrinovic, S., N. Menager, N. Goh, and P. Mastroeni, 2003 Characterization and development of T-Cell immune responses in B-cell-deficient (Igh-6(-/-)) mice with Salmonella enterica serovar Typhimurium infection. Infect Immun 71 (12):6808–6819.

Uthman, S., C. Bar, V. Scheidt, S. Liu, S. ten Have et al., 2013 The amidation step of diphthamide biosynthesis in yeast requires DPH6, a gene identified through mining the DPH1-DPH5 interaction network. PLoS Genet 9 (2):e1003334.

van der Weyden, L., M.J. Arends, A.D. Campbell, T. Bald, H. Wardle-Jones et al., 2017a Genome-wide in vivo screen identifies novel host regulators of metastatic colonization. Nature 541 (7636):233–236.

van der Weyden, L., N.A. Karp, A. Swiatkowska, D.J. Adams, and A.O. Speak, 2017b Genome wide in vivo mouse screen data from studies to assess host regulation of metastatic colonisation. Sci Data 4:170129.

van der Weyden, L., A.O. Speak, A. Swiatkowska, S. Clare, A. Schejtman et al., 2018 Pulmonary metastatic colonisation and granulomas in NOX2-deficient mice. J Pathol 246 (3):300–310.

Wang, J., J. Qian, Y. Hu, X. Kong, H. Chen et al., 2014 ArhGAP30 promotes p53 acetylation and function in colorectal cancer. Nat Commun 5:4735.

Webb, T.R., S.H. Cross, L. McKie, R. Edgar, L. Vizor et al., 2008 Diphthamide modification of eEF2 requires a J-domain protein and is essential for normal development. J Cell Sci 121 (Pt 19):3140–3145.

Wilk, E., A.K. Pandey, S.R. Leist, B. Hatesuer, M. Preusse et al., 2015 RNAseq expression analysis of resistant and susceptible mice after influenza A virus infection identifies novel genes associated with virus replication and important for host resistance to infection. BMC Genomics 16:655.

Yokoi, N., Y. Fukata, A. Sekiya, T. Murakami, K. Kobayashi et al., 2016 Identification of PSD-95 Depalmitoylating Enzymes. J Neurosci 36 (24):6431–6444.

Yokota, Y., A. Mansouri, S. Mori, S. Sugawara, S. Adachi et al., 1999 Development of peripheral lymphoid organs and natural killer cells depends on the helix-loop-helix inhibitor Id2. Nature 397 (6721):702–706.

Yu, Y.R., L.R. You, Y.T. Yan, and C.M. Chen, 2014 Role of OVCA1/DPH1 in craniofacial abnormalities of Miller-Dieker syndrome. Hum Mol Genet 23 (21):5579–5596.

Zago, G., M. Muller, M. van den Heuvel, and P. Baas, 2016 New targeted treatments for non-small-cell lung cancer - role of nivolumab. Biologics 10:103–117.

Zhang, J., J.L. Wang, C.Y. Zhang, Y.F. Ma, R. Zhao et al., 2019 The prognostic role of FZD6 in esophageal squamous cell carcinoma patients. Clin Transl Oncol.

Zhang, W., Z. Zhang, Y. Zhang, and A.P. Naren, 2017 CFTR-NHERF2-LPA(2) Complex in the Airway and Gut Epithelia. Int J Mol Sci 18 (9).

Zhou, Z., H. Jiang, K. Tu, W. Yu, J. Zhang et al., 2019 ANKHD1 is required for SMYD3 to promote tumor metastasis in hepatocellular carcinoma. J Exp Clin Cancer Res 38 (1):18.

